# HighDimMixedModels.jl: Robust High Dimensional Mixed Models across Omics Data

**DOI:** 10.1101/2024.05.09.593305

**Authors:** Evan Gorstein, Rosa Aghdam, Claudia Solís-Lemus

## Abstract

High dimensional mixed-effect models are an increasingly important form of regression in modern biology, in which the number of variables often matches or exceeds the number of samples, which are collected in groups or clusters. The penalized likelihood approach to fitting these models relies on a coordinate gradient descent (CGD) algorithm that lacks guarantees of convergence to a global optimum. Here, we study empirically the behavior of the algorithm across a number of common study types in modern omics datatypes. In particular, we study the empirical performance of high dimensional mixed-effect models fit to data simulated to mimic the features of transcriptome, genome-wide association, and microbiome data. In addition, we study the performance of the model on real data from each of these study types. To facilitate these simulations, we implement the algorithm in an open source Julia package `HighDimMixedModels.jl`. We compare the performance of two commonly used penalties, namely LASSO and SCAD, within the `HighDimMixedModels.jl` framework. Our results demonstrate that the SCAD penalty consistently outperforms LASSO in terms of both variable selection and estimation accuracy across omics data. Through our comprehensive analysis, we illuminate the intricate relationship between algorithmic behavior, penalty selection, and dataset properties such as the correlation structure among features, providing valuable insights for researchers employing high dimensional mixed-effect models in biological investigations.

**Author Summary:** High dimensional mixed-effect models are increasingly indispensable in modern biology, particularly in omics studies, where the number of variables often equals or surpasses the number of samples, and data are collected in clusters or groups. In our research, we concentrate on the penalized likelihood approach to fitting these models, employing a coordinate gradient descent (CGD) algorithm. While CGD is a widely used optimization technique, its convergence to a global optimum lack guarantees, prompting our empirical investigation into its behavior across various study types common in modern omics datasets. Our study provides insights into the performance of high dimensional mixed-effect models fitted to data simulating transcriptome, genome-wide association, and microbiome datasets. Additionally, we evaluate the model’s performance on real datasets from each of these study types. To facilitate reproducibility and further research, we have implemented the algorithm in an open-source Julia package, `HighDimMixedModels.jl`. Notably, `HighDimMixedModels.jl` stands out as the first package capable of seamlessly handling various omics datasets without errors, offering a user-friendly solution for researchers across disciplines. While numerous software packages are available for implementing high dimensional mixed-effects models on omics data, there is currently no comprehensive review source summarizing all methods. We provide a table summarizing existing methods, available in the Supplementary Material.

## Introduction

High dimensional regressions, in which the number of variables matches or exceeds the number of samples, are the rule, rather than the exception, in modern omics. In the field of transcriptomics, for example, RNA-Seq technology allows for simultaneous measurement of expression levels of thousands of genes [1, 2]; in population genetics, genome-wide association studies include hundreds of thousands, if not millions, of single nucleotide polymorphisms (SNPs) [3, 4]; and in metagenomics, microbiome studies measure the abundance (e.g., via 16S rRNA sequencing) of hundreds of bacterial taxa in perhaps only a few dozen samples from the environment [5, 6]. In order to make the analyses of these data sets tractable, it is typically assumed that only a small fraction of the covariates have a non-negligible effect on the response of interest, often related to human, soil or plant health. Identifying or selecting these impactful variables from the large pool of measured markers thus becomes a critical component of the analysis [7].

In many of these omics studies, samples are collected in groups or clusters. For example, observations might be made at different sites, such that samples taken from the same location are more similar to each other, or, in longitudinal analyses, collected in different individuals over time [8, 9], such that observations from the same individual may exhibit correlation. In a regression context, a common and flexible framework for handling this sort of grouped data is mixed-effects modelling. In a mixed-effects model, in addition to estimating the effects of any measured covariates, each group present in the data is also assigned an unobserved effect drawn from a common distribution. The sharing of the same random effect among units belonging to the same group induces correlation between these units [10].

Unfortunately, the standard approaches to fitting mixed-effects models, both (restricted) maximum likelihood methods and expectation-maximization algorithms, break down in the presence of high dimensional data sets. For example, if the number of covariates exceeds the total sample size, it becomes possible to find fixed effect regression coefficients that interpolate the data, and the maximum likelihood estimation of the model thus entails setting all variance components to 0, effectively eliminating any inter-group variation [11].

One common approach to estimating the parameters of a high dimensional regression models without random effects is via a penalized likelihood [12]. In this framework, parameters are chosen to maximize a loss function that is the sum of the model likelihood and a regularizing term that penalizes the size of the parameters. Penalizing the *ℓ*_1_ norm of the regression coefficient vector in an estimation method known as the least absolute shrinkage and selection operator (LASSO) has proven to be an extremely popular approach due to (1) the ease of the corresponding numerical optimization problem and (2) the sparsity of its solutions, consequences of the geometry of the *ℓ*_1_ norm and its convexity [13]. The properties of the LASSO as an estimator and as a feature selector have been extensively studied and have inspired a literature generalizing the LASSO and other sparsity-inducing regularization techniques to new domains [14].

The extension of the penalized likelihood estimation approach to the mixed-effects model has been recently studied in a number of papers [15–17]. In an initial paper [15], Schelldorfer, Bühlmann, and van de Geer studied the rate of convergence of the global maximizer of the penalized likelihood with an *ℓ*_1_ penalty and proposed a coordinate gradient descent (CGD) algorithm [18] for arriving at a local optimum of this objective function. Due to the non-convexity of the problem, the local optimum at which the algorithm converges is not guaranteed to coincide with the global optimum about which we have theoretical guarantees. Ghosh and Thorensen [16] subsequently studied the theoretical properties of regularized estimation of mixed-effects parameters under general, non-convex penalties, including the smoothly clipped absolute deviation (SCAD) penalty, which is able to produce sparsity similar to the LASSO while eliminating some of its bias in estimating non-zero parameters. Their analysis, however, is subject to the same qualification: even under this better behaved penalty, any theoretical guarantees apply only to the global optimizer and not necessarily to the local one arrived at through the descent algorithm.

Both [15] and [16] carry out simulations to study the performance of their estimation procedure under a single well behaved design matrix with normally distributed entries. Design matrices, however, vary in their structure across different omics studies, and the robustness of the CGD algorithm for fitting high dimensional mixed-effect models to these differences in design has not been studied empirically. In this paper, we set out to investigate the statistical performance of the numerical algorithm for fitting high dimensional mixed-effect models by way of simulation. In particular, we simulate design matrices that mimic common biological studies in the three settings mentioned above: gene expression, genome-wide association studies (GWAS), and microbiome studies. We then simulate responses according to the mixed-effects model and study the performance of the CGD algorithm with a LASSO penalty and with a SCAD penalty.

To facilitate this simulation study, we implement the CGD algorithm for fitting high dimensional mixed-effects models (HighDimMM) in a Julia package `HighDimMixedModels.jl` available at https://github.com/solislemuslab/HighDimMixedModels.jl. Our software mimics the code in R packages `lmmlasso` and `splmm` ([19, 20]), but includes the SCAD as the default penalty, which is not available in `lmmlasso` and whose implementation in `splmm` incorrectly updates the penalized fixed effects in the CGD algorithm. In addition, it corrects an error in the original code published in [16] that prevented zeroed coefficients from being further updated in the course of the algorithm.

While numerous software packages are available for implementing high dimensional mixed-effects models on omics data, there is currently no comprehensive review source summarizing all methods. We provide a table summarizing existing methods, available in the Supplementary Material [16, 17, 19, 21–50]. It includes paper names, package names, associated data sets, and crucial methodological details, such as whether the method involves a mixed-effects model with penalties and the application of penalties to random, fixed, or both effects. This comprehensive review aims to offer a detailed overview of available methods in this field. Our software distinguishes itself as one of the few (if not the only) that has been tested across omics data types. Indeed, most software has been created with a specific type of omics data in mind (e.g. gene expression or GWAS data), and the performance to other types of omics data is often not transferable. Our implementation works seamlessly and robustly across high dimensional omics data types, as shown in the simulations in this paper.

In the remainder of the article, we first present the formal high dimensional mixed-effects model, as well as the CGD algorithm for fitting it with LASSO and SCAD penalties, as found in [15] and [16]. Following this review, in Sections Simulation Studies and Applications to Real Omics Data, we describe, for each of the three omics study types, our strategy for simulating data and provide an example of a real data set that has appeared in published research. In Section Results, we detail the results of fitting high dimensional mixed-effects models with CGD to the simulated data and to the real data sets from each study type. We comment on the variation in the procedure’s ability to recover the impactful markers and accurately estimate their effects, across settings such as dimensionality, design matrix structure, and random effect structures. In Section Discussion, we conclude with some discussion of the major takeaways from our investigation.

## Results

### Simulation Studies

We fit models to each of the 100 data sets generated under each simulation setting in each study type in order to obtain Monte Carlo estimates of estimation performance under each setting. In each setting, we fit a well-specified model (i.e. we parameterized the covariance matrix for the random effects to include the true data generating random effects covariance matrix), and for each data set, the estimation of the model was done according to the procedure detailed in Section Computational Algorithm. Namely, across a grid of values of the regularization parameters *λ*, we attempted to minimize Equation (2), with penalty given by either the LASSO or SCAD, through the proposed CGD algorithm. We then chose the model that minimized the BIC across all the values of *λ*. In practice, we found that the local minimum to which the algorithm converged was quite sensitive to the choice of *λ*. With *λ* too low, the model converged to an interpolating solution, whereas setting *λ* too high resulted in all penalized regression coefficients being forced to zero. The range of *λ* that resulted in fits that were in between these two extremes could, for some data sets, be quite limited. We provide the grid of *λ*s that we searched over for the gene expression simulations in Table S3, with details on the grids for the other two simulation study types, GWAS and microbiome, in the caption.

We define the “final model fit” for each data set to be the fitted model that minimizes the BIC across the explored grid of *λ*s. In the following plots, we display the distribution of different statistics obtained from this final model fit across the data sets generated under a particular setting. In evaluating the final model fit’s performance, we focus primarily on the selection of the correct non-zero coefficients and on the accuracy of estimation for coefficients that are correctly selected. We first discuss the gene expression study type, then GWAS, and finally microbiome.

#### Gene Expression

##### Variable Selection

To evaluate the ability of the algorithm to perform accurate variable selection, we view each fitted model as performing a binary classification of the regression coefficients. In this context, we refer to coefficients that have been set to zero in the fitted model as “negatives” and coefficients estimated non-zero as “positives”. A true positive is a coefficient correctly estimated non-zero, meaning that its ground-truth value is also non-zero. A false positive is a coefficient incorrectly estimated non-zero. We define the false positive rate (FPR) for a given model fit to be the proportion of ground-truth zero regression coefficients that are mistakenly selected (i.e. estimated non-zero):

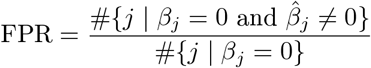

In Figure 1A, each box plot displays the distribution across data sets of false positive rates of the final model fit upon employing the LASSO (orange) or SCAD (blue) penalty to each of the the data sets simulated under a given setting. This figure covers all settings with *q* = 3 random effects (settings 7-12 in Table 3); we provide the analogous figures for all other settings (*q* ≠ 3) in Figure S1.

**Figure 1:**
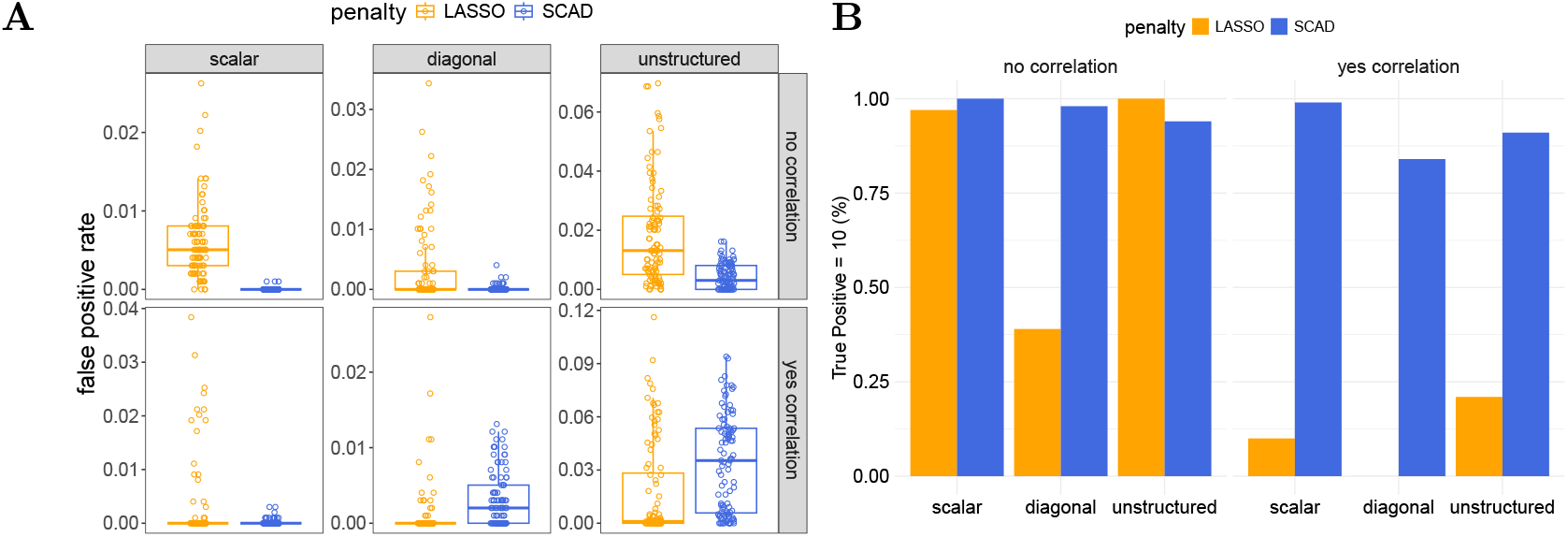
Variable selection of final model fits to all gene expression data sets simulated with *q* = 3 random effects. (settings 7-12 in Table 3). In panel A, the columns identify different random effect covariance (Ψ_*θ*_) structures, while the row faceting indicates the presence of inter-feature correlation. “no correlation” = no inter-feature correlation, “yes correlation” = yes inter-feature correlation. The Y-axis corresponds to false positive rate. In Panel B, the height of each bar indicates the proportion of data sets in which the model recovers all 10 true non-zero coefficients. For complete results across different ranges of true positive rates, refer to Table S4.

In all but three settings, the median number of false positives from using the SCAD penalty was zero. The most challenging settings for the SCAD penalty were those with correlated features and three random effects following an unstructured or diagonal covariance matrix (settings 10 and 12). In every other setting, the LASSO-based estimator included false positives more frequently and in higher number. The LASSO-based estimator struggled most in the setting 14 with five random effects and correlated features (Figure S1C), where its median false positive rate was 27/990 and maximum as high as 184/990 = 18.6%. Some of these results may be artifacts of searching over a finer grid of penalty hyperparameters for the models with greater covariance complexity (see Table S3).

The bar graphs in Figure 1B represent the proportion of data sets in which the final model fit had perfect recall, i.e. recovered all 10 true non-zero coefficients (analogous figures for settings with *q* ≠ 3 in Figure S1B and D). Estimation with the SCAD penalty recovered all 10 non-zero coefficients in the vast majority of data sets. We additionally see from comparing Figure 1A and 1B that in the two settings in which the LASSO penalty led to a lower median FPR than the SCAD penalty, this was at the cost of having much worse recall.

##### Estimation

The LASSO penalty is known to bias its estimates of non-zero parameters towards zero [51]. In our gene expression simulations, we find that this LASSO-penalty-based bias can lead to bias in the estimates of even the coefficients that are not penalized (in the context of mixed-effect models, such unpenalized coefficients are often the coefficients on features for which there are random effects, i.e. columns of *Z*_*i*_). In Figure 2, we have displayed the distribution of estimates of one of the penalized regression coefficients, *β*_4_, and one of the unpenalized regression coefficients, *β*_3_, across all gene expression data sets simulated with *q* = 3 random effect (settings 7-12 in Table 3) with analogous plots for all other settings given in Figure S2. Specifically in the settings with inter-feature correlation, the LASSO-based estimates of the non-penalized *β*_3_ form two clusters, neither of which are centered on the true parameter *β*_3_ = 4, but which are biased to varying degrees. This bias results from the penalization applied to *β*_4_, as *β*_3_ and *β*_4_ are the coefficients on adjacent columns in the design matrix, which have correlation *ρ* = .6 in these settings. The upward bias in the estimator for *β*_3_ is compensating for the downward bias in the estimate of *β*_4_, with the required compensation largest when *β*_4_ is completely excluded from the selected variables, resulting in the observed clustering in the estimates of *β*_3_. This is additionally visualized in Figure S3, a scatter plot of the LASSO-based estimates of these two coefficients in the presence of inter-feature correlation. Because the estimation of *β*_4_ under the SCAD penalty is much more accurate – it is correctly included in the active set in the vast majority of cases and estimated without bias when it is – its bias in the estimation of *β*_3_ (in the settings with inter-feature correlation) is much less acute than the LASSO’s. In general, the SCAD estimates for all penalized coefficients were unbiased in all settings, conditional on their selection (blue box plots are centered on the true parameter value which is represented by a solid black line). The standard deviation of the SCAD estimates, conditional on their selection, ranged from 0.043 in the easiest setting with 1 random effect and no inter-feature correlation, to 0.12 in the setting with 3 random effects following an unstructured covariance matrix and no inter-feature correlation (Table 1).

**Table 1:** Effect estimation for gene expression data. Statistics summarizing the Monte Carlo distribution of the estimates of *β*_8_ when using SCAD penalty in each of the settings with 10 non-zero regression coefficients (settings 5-14 from Table 3 where *N* = 250 and *p* = 1000), conditional on the estimate being non-zero. The true value of the parameter being estimated was -3. The column 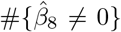 indicates the number of data sets, out of 100, for which we correctly estimate *β*_8_ to be non-zero (and is thus the sample size with which these summary statistics are calculated).

| $q$ | $\Psi_\theta$ | Correlated Features | $\#\{\hat{\beta}_8 \neq 0\}$ | $E[\hat{\beta}_8 \vert \hat{\beta}_8 \neq 0]$ | $SD[\hat{\beta}_8 \vert \hat{\beta}_8 \neq 0]$ |
| --- | --- | --- | --- | --- | --- |
| 1 | scalar | No | 100 | -3.00 | 0.04 |
| 1 | scalar | Yes | 100 | -3.01 | 0.08 |
| 3 | scalar | No | 100 | -3.00 | 0.06 |
| 3 | scalar | Yes | 100 | -2.99 | 0.08 |
| 5 | scalar | No | 99 | -3.00 | 0.08 |
| 5 | scalar | Yes | 100 | -2.99 | 0.11 |
| 3 | diagonal | No | 98 | -2.99 | 0.06 |
| 3 | diagonal | Yes | 84 | -3.01 | 0.09 |
| 3 | unstructured | No | 95 | -3.00 | 0.07 |
| 3 | unstructured | Yes | 91 | -2.99 | 0.12 |

**Figure 2:**
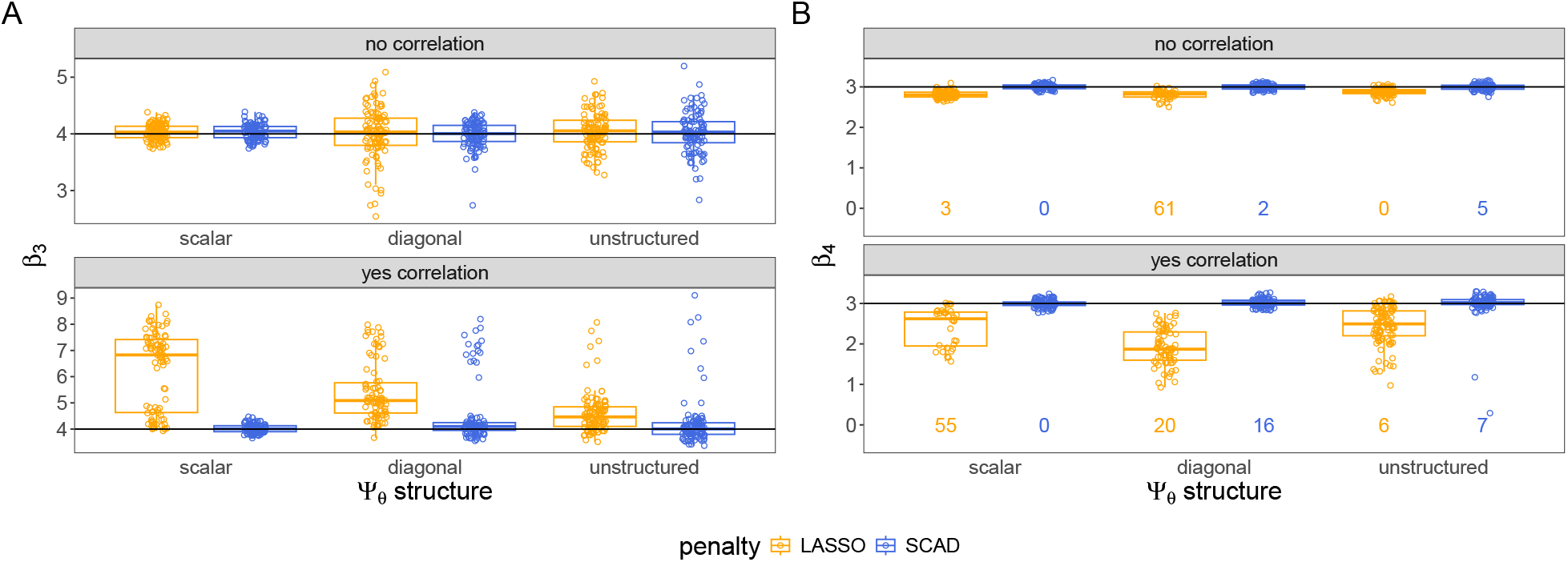
Effect estimation for gene expression data. Distributions of estimates of an unpenalized regression coefficient (A) and of a penalized regression coefficient (B) across all gene expression data simulated with *q* = 3 random effects (settings 7-12 in Table 3). Numbers at the bottom of the plotting windows in B indicate the number of data sets for which the coefficient was incorrectly estimated 0. The box plots represent the distribution of the estimator across all other data sets (i.e. all data sets in which the estimate was non-zero). The true parameter value is represented by a solid black line.

In most applications, the estimation of variance components is of secondary importance relative to the estimation of fixed effect parameters. Figure S4 shows our estimates of elements of scalar, diagonal, and unstructured random effect covariance matrices, respectively, in the settings with 10 non-zero regression coefficients. Diagonal entries of these random effects covariance matrices were estimated zero when the estimated fixed effect vector included many false positives, as these additional non-zero coefficients proved to be an alternative means to fit the variability driven by the random effects. The LASSO estimates included false positives more frequently, and in these models with many false positives, the random effect variance components were frequently set to zero.

#### GWAS

For the GWAS simulations, we fit a model with SCAD penalty and a model with LASSO penalty to each combination of design matrix and response vector.

##### Variable Selection

We illustrate the performance of our estimators at the task of variable selection in the GWAS setting for the data sets with *q* = 3 random effects in Figure 3 (results in simulations with *q* = 1, 5 are shown in Figure S5). Both SCAD and LASSO estimators were able to recover the 10 true non-zeros with similar consistency, only occasionally missing one or more of them (Figure 3B). Panel A of figures 3 and S5 illustrate, however, that the LASSO penalty led to more false positives than the SCAD penalty in each setting.

**Figure 3:**
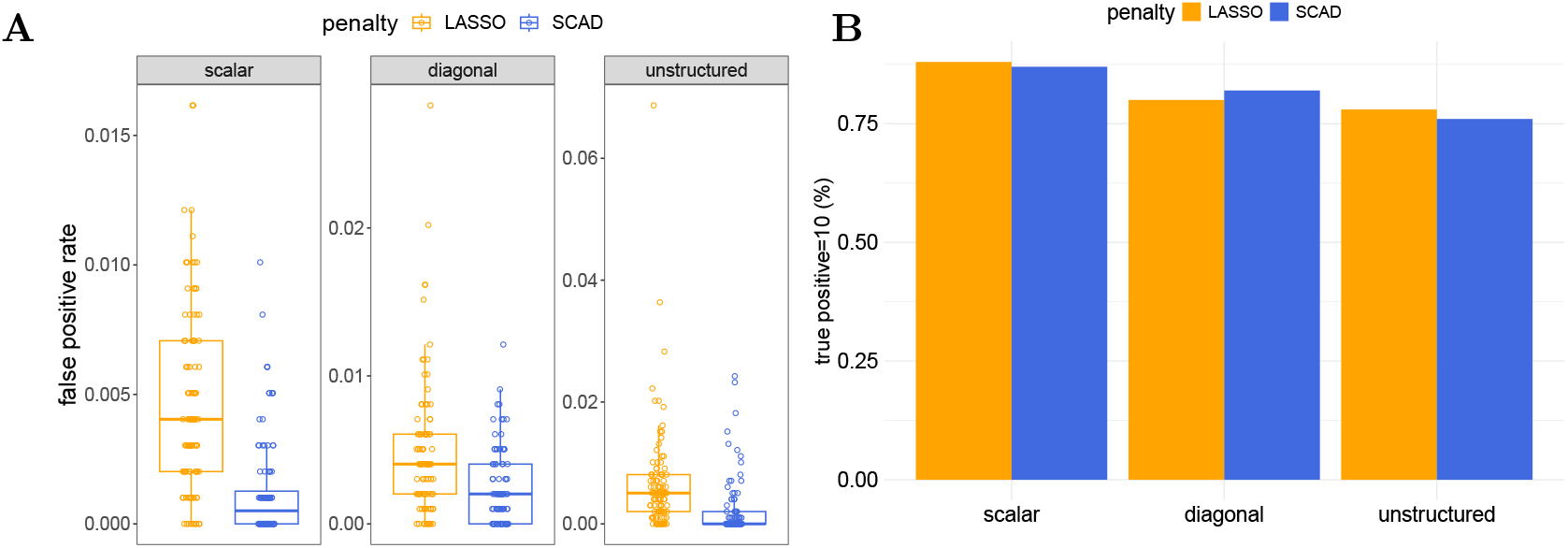
Variable selection for GWAS data. Performance at task of variable selection on GWAS data simulated with *q* = 3 random effects. In Panel A, Y-axis corresponds to false positive rate and the facets identify different random-effect covariance matrix structures (Ψ_*θ*_). In panel B, the height of each bars indicates the proportion of simulated data sets in which the model identifies all 10 true non-zero coefficients. Colors again differentiate the penalty used to fit the model. For complete results across different ranges of true positive rates, refer to Table S5.

##### Estimation

We display the distribution of the estimates of one unpenalized regression coefficient, *β*_3_, and one penalized regression coefficient, *β*_8_, from the GWAS simulations with *q* = 3 in Figure S6 (estimates for settings with *q* = 1 or 5 in Figure S7). In the GWAS simulations, because the features (counts of minor alleles at different locations in the genome) are uncorrelated, we observed no bias in the estimation of non-penalized coefficients under the LASSO (and as usual, the SCAD-based estimators were also unbiased). On the other hand, the bias towards zero in estimates of penalized coefficients under the LASSO penalty remained a problem in the GWAS simulations (Figure S6B).

In terms of the variance components, we observe that the estimates were significantly less accurate than the estimates in the gene expression simulations (Figure S8). Of course, we can attribute this to the reduction in the number of clusters from *g* = 50 in the gene expression settings 3-14 from Table 3 to *g* = 10 in the GWAS simulations. In general and as previously mentioned, variance components are typically of secondary interest to the fixed effect estimates in -omics studies. Nonetheless, it is worth noting that their estimation improves with the number of clusters, as one would expect from experience with low dimensional mixed-effect models.

#### Microbiome Data

Finally, we describe our results in the microbiome setting. Recall that for these simulations, we simulated OTU count matrices under six different assumed latent OTU network structures, as we were interested in the downstream impact of the count matrices’ correlation structure on model performance.

We fit models to each microbiome data set, consisting of a log-ratio transformed microbiome design matrix and response vector. Having observed the superior performance of SCAD relative to LASSO penalization in the other two settings, we focused on only the SCAD penalty in the microbiome setting. We refrained from penalizing the coefficient on the variable measured at the group level (and as usual, did not apply a penalty to coefficients that had corresponding random effects). With the addition of the coefficient on this group-level variable, there were thus 11 non-zero ground-truth regression coefficients for these simulations, which we hoped to recover and accurately estimate.

##### Variable Selection

Since the “band” network structure produced design matrices that proved to be the most challenging to the model, whereas the “scale-free” structure led to the best results, we display model performance from these two network structures in the main text to communicate an accurate sense for the range of results. Figure 4 shows the performance of the estimation procedure at the task of variable selection for these OTU structures for simulations with *q* = 3 random effects.

**Figure 4:**
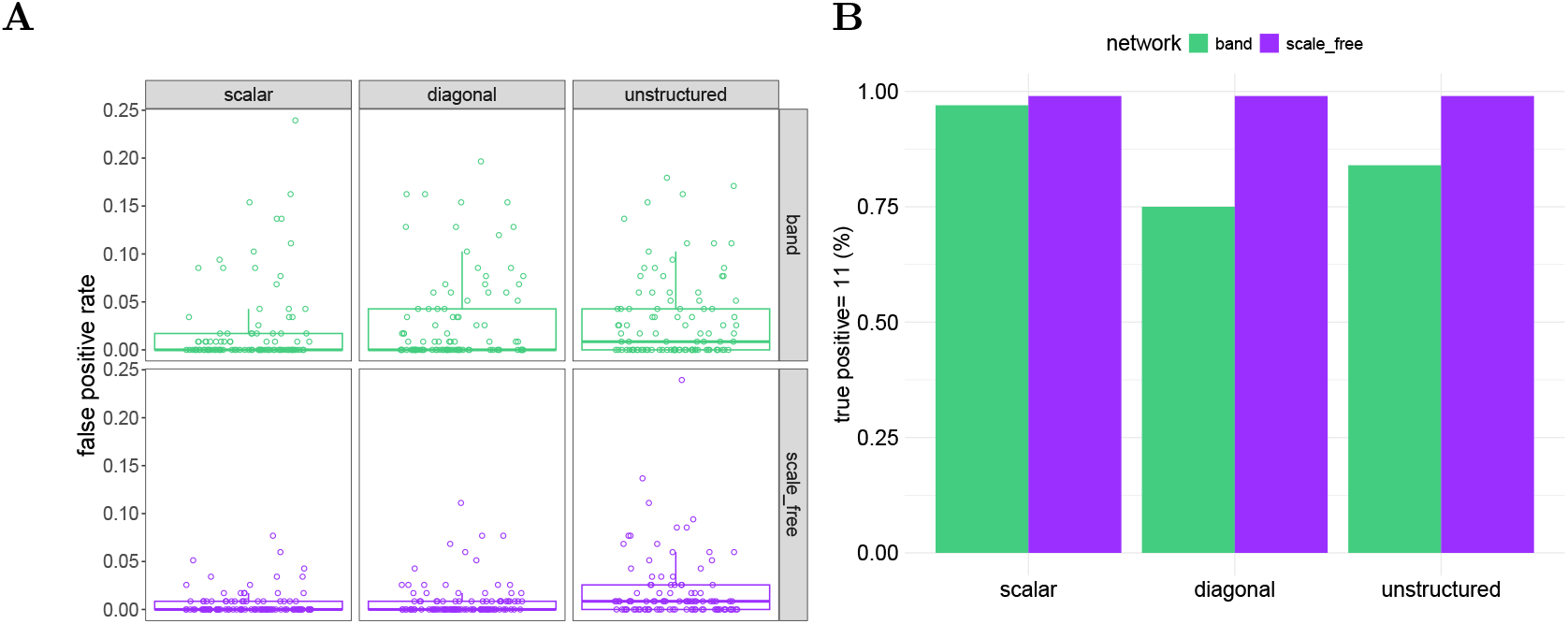
Variable selection for microbiome data. Performance at task of variable selection on microbiome data simulated under a “scale-free” or “band” OTU network structure and with response generated with *q* = 3 random effects. In panel A, the columns identify different random effect covariance (Ψ_*θ*_) structures, while the row faceting (and color) indicates the OTU correlation-structure. The Y-axis corresponds to false positive rate. In panel B, the height of each bars indicates the proportion of simulated data sets in which the model identifies all 11 true non-zero coefficients. For complete results across different ranges of true positive rates, refer to Table S6.

We recovered all eleven regression coefficients in almost every case when design matrices were generated from a scale-free OTU network, and when the design matrices was generated from a band OTU network but the random effect covariance matrix was scalar. When the design matrices were generated from a band OTU network and the random effect covariance matrix was more complex (diagonal or unstructured), the recall of the model suffered, and in particular, we frequently converged to a solution with all penalized coefficients set to zero (resulting in four true positives from the non-penalized coefficients) (see Table S6). The false positive rates were comparable across settings, with slightly higher false positive rates in the settings that had worse recall, as the model compensated for setting true effects to zero.

The variable selection results for the scale-free and band network structures with *q* = 1 and *q* = 5 are displayed in Figure S9. We see that the model struggled when there were *q* = 5 random effects and a band design matrix, frequently setting all penalized coefficients set to zero. In the other four OTU network structures, the variable selections results were on par with the results on the scale-free results or slightly worse (Figure S10). Notably, the algorithm did not converge as frequently to a solution with all penalized coefficients set to zero under these other four OTU-OTU covariances as it did with the band OTU-OTU covariance.

##### Estimation

The fixed effect coefficient on the variable measured at the group level was estimated without bias in the OTU simulations (shown for *q* = 3, band and scale-free network structures in Figure S11, and other settings in Figures S12 and S13). Since we used a SCAD penalty, we also estimated all coefficients of covariates measured at the individual level without bias, as usual. There was more variance in the estimates of the coefficient on the group-level variable, which is not surprising, given that there are far fewer groups than individuals. The data sets that led to extreme outliers in our estimates of this coefficient were the ones in which the estimate of penalized coefficients were mistakenly set to zero, frequently data sets with a band OTU covariance structure and three or five random effects.

### Application to Real Omics Data

We now turn to our real data examples, presenting them in the same order as our simulations.

#### Bacterial Gene Expression and Riboflavin Production

We adopt the data-driven strategy found in [15] to select columns to receive random effects and fit a model that includes the selected random effects (and, in contrast to previous models, no random intercept) with a diagonal covariance structure. As in [15], the strategy to identify random effects leads to the assignment of random slopes to two genes, *YFJD* and *YTOI*. We fit a model that included random slopes for these two genes (and in contrast to previous models, no random intercept) with a diagonal covariance and with *λ* set to 45 based on the BIC. This final model selects the 17 genes listed in Table 2 as potentially impacting the riboflavin production rate. In order to comprehensively evaluate the list of genes resulting from our analysis, which initially identified 17 genes, we applied the generalized linear mixed model method, as proposed in [52], to the Riboflavin dataset, similar to the approach outlined in Buhlmann et al. [7]. This analysis yielded a set of 28 significant genes. Upon comparison, we identified intersecting genes *YDDK, YURQ, LYSC*, and *YXLD* between these two gene lists. Additionally, following the methodology described in [7], we employed high dimensional mixed-effect models with the LASSO and SCAD penalties, denoted as lmmlasso and lmmSCAD, respectively, on the same dataset. By treating genes *YFJD* and *YTOI* as random effects and setting lambda values at 45 for LASSO and 25 for SCAD with a diagonal covariance, we identified 28 important genes for lmmlasso, of which 3 (*TUAH, YXLD, YDDK*) were found to be common with our gene list. For lmmSCAD, the analysis yielded 5 intersecting genes (*LYSC, TUAH, YURQ, YXLD, YDDK*). The complete gene lists are provided in Table S7, with intersecting genes highlighted in bold.

**Table 2:** GWAS hits on riboflavin data. Genes with non-zero estimated effect on riboflavin production using our HighDimMM model

| Gene | Estimated-effect | Gene | Estimated-effect |
| --- | --- | --- | --- |
| *YURQ* | 1.427 | *YCGP* | -0.103 |
| *YFKE* | 1.073 | *YDBM* | -0.103 |
| *YTOI* | 1.019 | *YTXM* | -0.115 |
| *YFJD* | 0.721 | *UVRB* | -0.176 |
| *YLMA* | -0.003 | *YNEI* | -0.235 |
| *YUSY* | -0.005 | *LYSC* | -0.383 |
| *METK* | -0.025 | *YXLD* | -0.526 |
| *YUBB* | -0.055 | *TUAH* | -1.343 |
| *YDDK* | -0.1 |  |  |

The estimated error variance for this final model was 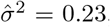, on par with the error variance in [15]. Finally, although we allowed for two random effects in the final model, one of these error variances is set to zero over the course of the computational algorithm, leaving only the gene *YFJD* with an effect varying across bacterial cultures (with estimated variance 0.38). In this model, as in all models we fit to real data sets in the coming sections, we used the SCAD penalty, given its superior performance in the simulation studies.

#### Mouse GWAS Study for Body Mass Index

We fit high dimensional mixed-effect models with the SCAD penalty to the mouse data set, searching over three different values of the regularizing hyperparameter *λ*. As in all previous models, increasing *λ* resulted in fewer selected loci, and the estimates from all three are shown in Figure 5. The fit that minimized the BIC was obtained with the largest penalty and included only 7 SNPs. The predictive accuracy of this final model was on par with the Bayesian model from [53], as the estimated error variances of the two models were 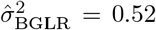 and 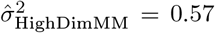, respectively. Because the Bayesian model does not apply a sparsity-inducing penalty, however, it includes all SNPs in the model and is therefore less interpretable than our model fit with the SCAD penalty. We validated the features selected by our model by checking that all 7 SNPs with estimated non-zero coefficients from our model are among those with the top 20 largest coefficient magnitudes in the Bayesian model. Based on Figures 5 and S14, our model is able to reduce the error variance as much as the Bayesian model while using far fewer features by 1) assigning larger effect sizes (coefficient magnitudes) to the selected features and 2) making greater use of the cage effects, as seen by comparing the observed versus fitted plots with and without the inclusion of the random intercepts in Figure S14.

**Figure 5:**
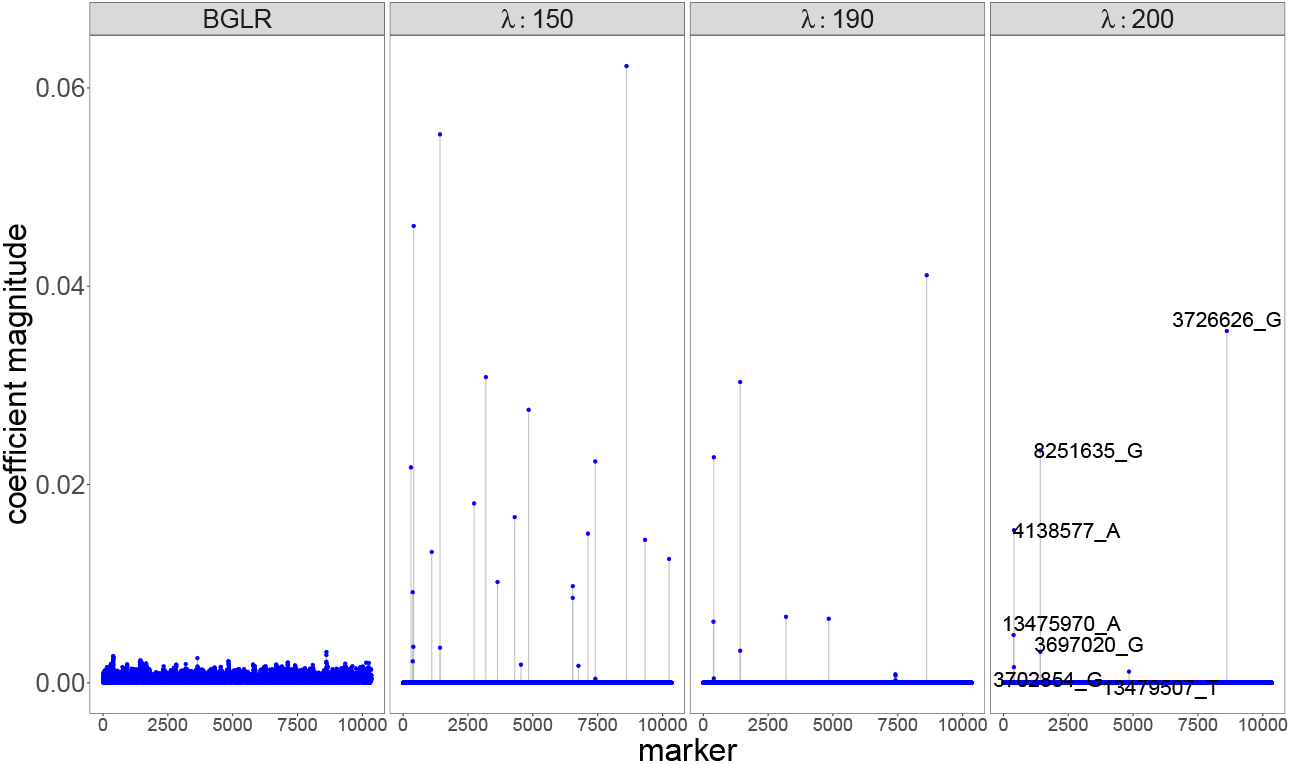
Absolute values of coefficient estimates from HighDimMM using a SCAD penalty with different values of *λ* and from a Bayesian model fit with BGLR which does not use a sparsity-inducing penalty and thus includes all SNPs in the model. The value of *λ* is given in the strips above the plots–the plot on the far right is from BGLR. For our model, the fit that minimized the BIC was obtained with *λ* = 200 and included only 7 SNPs, which are labelled in the plot.

We additionally compared our results to the analysis of the same dataset performed in [50]. Their model selects 13 SNPs as associated with BMI at a false discovery rate (FDR) of 0.05, in addition to an effect of gender. Of these 13 SNPs, 2 are also selected in our model fit with *λ* = 150, but none are selected in our model fit with *λ* = 190 and *λ* = 200. The two SNPs that overlap with our model fit with *λ* = 150, which selected 25 SNPs in total, are *rs3023058* and *rs6185805*. These two SNPs are located on the genes *Srrm3* and *Mtcl1*, respectively.

As previously mentioned, we differ slightly from the model specification in [50] in our choice to include mouse litter, a categorical variable that is shared by all mice in the same cage, as a predictor in the model. Our model produced (unpenalized) estimates of 0.4 and 0.5 (which are substantial effect sizes given that the largest SNP coefficient is an order of magnitude lower at 0.04) for the coefficients on litters 7 and 8 (and negligible coefficients for all other litters). However, given that these were the smallest litters with sample sizes of only 14 and 4, these effects are not necessarily statistically significant, and the inclusion of litter as a predictor likely does not explain the large discrepancy in identified SNPs by our approach and that of [50]. More likely, the lack of overlap in selected SNPs points to a fundamental differences between the likelihood-based approach to fitting the model and the quasi-likelihood approach proposed in [50].

#### Human Gut Microbiome Data across Age and Geography

When we fit the model to the library-sum-scaled OTU data, we find that the algorithm converges to a solution with only 27/1362 non-zero regression coefficients. In the context of the data, we’ve identified these 27 OTUs as being conditionally associated with age controlling for country and for all other OTUs (Table S8). The absolute values of the non-zero regression coefficients ranged from 0.64 to 1424.94. Recalling that the response is the square root of age and the design matrix consists of OTU percent compositions, the largest of these coefficient represents a non-trivial shift in the distribution of age: for every increase of 1% in the microbial composition of the OTU, the expected value of the conditional distribution of the square root of age increased by 1.4. The model *R*^2^ was 63%, on par with the *R*^2^ of models for the same data presented in [28].

## Discussion

In this work, we have provided the first working implementation of the CGD algorithm for fitting high dimensional mixed-effect models with the SCAD penalty, and we have tested the statistical performance of the algorithm across three canonical data types in modern biology. We highlight here that the CGD algorithm fit with the SCAD penalty was found to yield accurate feature selection and estimation even for unusual design matrices such as those generated by applying a log-ratio-transformation to microbiome count data, and even when including variables measured at the group level in the design matrix. For microbiome studies, these variables often represent features of the environment in which the distinct clusters of microbiome samples are collected, such as temperature or soil quality.

Previous implementations of the CGD algorithm for high dimensional mixed-effect models have 1) failed to implement the SCAD penalty all together (`lmmlasso`), 2) failed to implement the correct update of the penalized coefficients under the SCAD penalty (`splmm`), or 3) failed to correctly implement the active set strategy, resulting in an algorithm in which any regression coefficient that is set to zero at a given iteration is never further updated (`lmmsSCAD`). In contrast, our algorithm correctly implements both the algorithmic updates for the SCAD penalty, as well as the meta-algorithmic active set strategy. In our simulation studies, we found that the SCAD penalty provided substantial improvements in terms of both variable selection and estimation accuracy relative to the LASSO penalty. Thus, our implementation of the algorithm with the SCAD penalty in a working and easy-to-use Julia package should be useful to biologists working across genomics who wish to fit these models to analyze their clustered, high dimensional data.

At the same time, several limitations of these models have become clear to us over the course of our simulation studies. First, the fitting of these models requires the choice of the regularization hyperparameter *λ*. Choosing a *λ* that is too small results in a model that includes many predictors and interpolates the data; meanwhile, a *λ* that is too large results in a model in which all penalized coefficients are set to zero. While this is a characteristic of all penalized likelihood estimation strategies, it poses a particular challenge for models in which fitting the model for a particular choice of *λ* is relatively computationally expensive, such as coordinate descent algorithms (even with the active set strategy), since the entire model selection process requires repeating this computationally expensive procedure across a grid of *λ*s. For this reason, a *pathwise coordinate descent* approach, in which the algorithm for each *λ* is initialized with the solution for the previous *λ* in the path, has been a popular approach to efficiently fit LASSO models without random effects [54]. It is worth exploring whether this approach could be extended to the fitting of high dimensional models with random effects, using either the LASSO or SCAD penalty.

The second major limitation of this approach to analyzing high dimensional, clustered data is the current lack of tools for assessing the degree of certainty associated with the selected features. Buhlmann and et al. [7] present a variety of strategies for quantifying feature selection uncertainty in high dimensional biological studies. Although these strategies do not explicitly account for random effects, many of them may still prove useful in the mixed-effect context. One complication is that many of these strategies rely on some form of bootstrapping and in a clustered data context, bootstrapping is less straightforward. Nonetheless, in future work, we hope to adapt some of these strategies in order to quantify the uncertainty and operating characteristics of the features selected by the penalized mixed-effect model. Post-selection inference for mixed-effect models is an area of ongoing research.

Finally, we note that the model and likelihood presented in this paper are that of a continuous, Gaussian response, and we have not provided any advice for biologists working with other types of responses, such as binary or count data. There are existing algorithms and methods (see [32], [55]) that can be used when the distribution of the response is a member of an exponential family – that is, for fitting generalized linear mixed models in the high dimensional setting – that we have not studied in the present work. Obviously, many biological responses are not continuous and Gaussian, and these methods are required for these cases. It is important to have an understanding of the operating characteristics of these methods – in particular, the reliability of their selection of predictors – and a simulation study similar in scope to the one we have conducted in this paper, with dataset examples drawn from the biological study types we have identified but with differently distributed responses, would be valuable towards this end.

## Materials and Methods

### Model Formulation and Penalties

Let *g* denote the number of clusters in our data. In this paper, we consider the linear mixed-effects model, where the vector of responses *y*_*i*_ in each cluster *i* = 1, …, *g* is generated according to

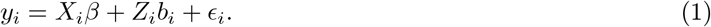

We assume that we are performing variable selection of the fixed effects rather than the random effects so that *β* ∈ ℝ^*p*^ is a high dimensional vector of fixed effects, and the 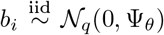 are low dimensional vectors of random effects (*q << p*). Letting *n*_*i*_ indicate the number of observation in cluster 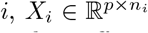 and 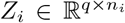 are cluster specific design matrices, with the latter corresponding to the random effects in the model, such that the variables represented as columns of *Z*_*i*_ are taken to vary across the clusters in their effects on the response. These variables are typically a subset of the columns in *X*_*i*_, so that although the random effects are drawn from a mean zero distribution, the average effect across the clusters may be estimated non-zero as a fixed effect. Finally, for each cluster 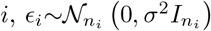 is a vector containing the error terms for each response in the cluster. The *ϵ*_*i*_ are assumed mutually independent of each other and of the random effects.

The covariance matrix Ψ_*θ*_ of the random effects is a symmetric positive definite matrix that is parameterized by *θ* ∈ ℝ^*q*\*^ . We consider three distinct models in which Ψ is, in turn, a scalar (*q*^*^ = 1), diagonal (*q*^*^ = *q*), and arbitrary symmetric positive definite (*q*^*^ = *q*(*q* + 1)*/*2) matrix. In each case, *θ* is the lower-triangular Cholesky factor of Ψ_*θ*_ and, as such, can be optimized without any constraints.

The marginal distribution of *y*_*i*_ in this model is *𝒩* (*X*_*i*_*β, V*_*i*_(*θ, σ*^2^)), where 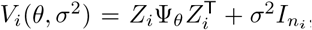, and the log-likelihood of the full set of parameters *ϕ* = (*β, θ, σ*^2^) is thus

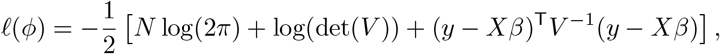

where *y* ∈ ℝ^*N*^ and *X* ∈ ℝ^*N×p*^ are obtained by vertical stacking, *V* ∈ ℝ^*N×N*^ by diagonal stacking, and *N* denotes the total sample size obtained by summing the cluster sizes, 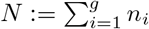. We define the maximum penalized likelihood estimators to be the minimizers of the loss function

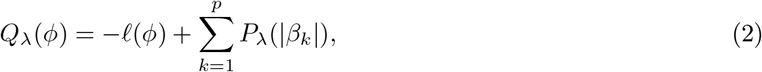

where *λ* is a hyperparameter that governs the severity of penalization and must be selected. In the initial study of this model, an *ℓ*_1_ penalty *P*_*λ*_(|*β*_*k*_|) = *λ*|*β*_*k*_|, was proposed [15]. As an alternative, the SCAD penalty, defined through its derivative by

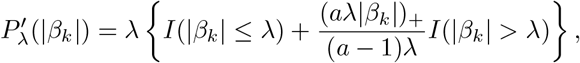

was proposed for penalized regression originally in the classic low-dimensional set-up [51], subsequently in the high dimensional regime [56], and finally for high dimensional mixed-effects models in [16].

### Computational Algorithm

A CGD algorithm for minimizing the loss function in Equation (2) was originally proposed in [15] and is implemented in our package. The algorithm is originally introduced in [18] but tailored to the penalized mixed-effect loss function in [15] for the LASSO penalty and in [16] for the SCAD penalty. At every iteration *k* of the algorithm, we cycle through indices 1, 2, … *p* + *q*^*^ + 1, and update each component as follows:

1. For the components of *β*, i.e. *ϕ*_*j*_, *j* ∈ *{*1, …, *p}*, we update the estimate 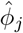 by minimizing a quadratic approximation of the loss function centered at the current estimates while all other components are kept fixed at their current values.
2. For the variance-covariance parameters, i.e. *ϕ*_*j*_, *j* ∈ *{p* + 1, …, *q*^*^ + 1*}*, we update the estimate 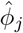 by simply maximizing the likelihood with respect to this component while all other components are kept fixed at their current values.

Steps 1 and 2 are repeated until the estimates of both fixed effects and variance-covariance components converge. Step 1 is computationally expensive, as it requires updating every component of a high dimensional vector. To considerably reduce the computation, following [15], we adopt an “active set” strategy [39, 52]: instead of updating every component of 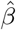 at each iteration, we update only those that are currently non-zero, with updates to the full set of components made only every *D* (*D* = 5 in our simulations) iterations. In addition, we update the full vector any time the convergence criterion is satisfied after an update of only the active set. Only when the convergence criterion is satisfied after a full update do we terminate the algorithm. This constitutes a single fit of our model for a given value of the regularization hyperparameter *λ*. As in [15, 16], we search over a grid of *λ*s, fitting a model with each one, and selecting as our final model the fit that minimizes the Bayesian information criterion 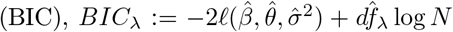, where the estimated degrees of freedom 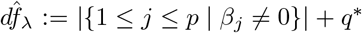 is the number of parameters estimated non-zero.

### Simulation Studies

We simulated three types of omics data sets – transcriptome profiles, GWAS data, and 16S microbiome data – in order to empirically study the performance of the CGD algorithm in fitting high dimensional mixed-effects models across these biologically meaningful scenarios. In each, the response was generated according to Equation (1), but the form of the design matrices *X*_*i*_ and *Z*_*i*_ varied considerably across the three studies so as to mimic their real-world counterparts. Within each study, we additionally varied aspects of the data such as dimensionality, sample size, and random effect structure to investigate their effect on performance. In the following sections, we detail the settings that we entertained within each of the three study types.

#### Gene Expression Simulations

To simulate gene expression design matrices, we draw a sample of *N* i.i.d. random vectors of length *p*−1 from a multivariate normal distribution. Viewing these vectors as (standardized) profiles of the transcritomes of the collected samples, they become the rows of our design matrix *X*; meanwhile, the first *q* − 1 components of these vectors become the rows of the matrices *Z*_*i*_ and thus receive random effect (*p* − 1 and *q* − 1 because the final columns are set as constants for estimating fixed and random intercepts, respectively). The multivariate normal distribution from which the rows of *X* are drawn has mean zero and a first order auto-regressive covariance matrix: Cov(*X*_*i,j*_, *X*_*i,j′*_) = *ρ*^|*j*−*j′*|^ for some choice of *ρ* ≥ 0. Thus, for non-zero *ρ*, columns of the design matrix that are closer to each other in index are more correlated. For the dimensionality *p* of the problem, we consider sizes of 500 and 1000, typical of the number of genes in a small scale transcriptomics study. We set the sample size to be *N* = 180 when *p* = 500 and *N* = 250 when *p* = 1000, creating a regime in which regularization is necessary for avoiding interpolation. We view each sample as belonging to a cluster: for the settings with a total sample size of *N* = 180, we viewed the samples as belonging to 30 different clusters, each of size 6, whereas for the settings with a total sample size of 250, we considered there to be 50 clusters, each of size 5.

After simulating the design matrices *X*_*i*_ and *Z*_*i*_ for each cluster *i*, we generated the response *y*_*i*_ according to Equation (1). In the settings with a sample size of *N* = 180, we chose only 5 (out of *p* = 500, corresponding to a sparsity of 5*/*500 = 1%) of the components of the fixed effect vector *β* to be non-zero and included only a random intercept, but no random slopes (*q* = 1). We refer to the set of indices of the non-zero regression coefficients *{*1 ≤ *j* ≤ *p* : *β*_*j*_ *≠* 0*}* as the “active set”. In the larger scale simulations in which *N* = 250 and *p* = 1000, we conducted a more extensive exploration of different combinations of fixed effect active set sizes and random effect structures. For the fixed effects, we considered the original active set, as well as the potential impact of doubling its size to match the sparsity of the smaller simulations (10*/*1000 = 5*/*500 = 1%). For the random effect structure, we varied the number of random slopes, investigating *q* ∈ *{*1, 3, 5*}*; the *q* = 1 case corresponds to a model with only random intercepts, while the other two cases include random slopes for two or four of the predictors, respectively. Furthermore, with *q* = 3, we simulated data under three different choices of the covariance matrix of the random intercept and slopes: a scalar matrix (multiple of the identity), a diagonal matrix, and a general symmetric positive semi-definite matrix. For *q* = 5, we considered only a scalar covariance matrix to manage the complexity of the model.

Table 3 summarizes each of the settings entertained in our gene expression simulations, and Table S1 details the values of the parameters *β*, Ψ_*θ*_, and *σ*^2^ in these simulations (Ψ_*θ*_ in table, *β* and *σ*^2^ in caption). Note that our setting 2 is identical to setting “H2” in [15] (with identical parameters). We go beyond [15] in our extensive exploration of other settings, focusing on the impact of dimensionality, sparsity, and random effect structure. In each of the fourteen settings under investigation, we simulated 100 data sets, consisting of matrices *X* and *Z*, response vector *y*, and a vector tracking the group membership of each observation.

**Table 3:** Gene expression simulation settings. Fourteen different settings under which we generated transciptomics data sets. Setting differed in sample size (*N*), number of clusters (*g*), number of fixed effects (*p*), number of random effects (*q*), correlation in auto-regressive covariance matrix (*ρ*), number of non-zero fixed effects (# effects) and random effects covariance (Ψ_*θ*_) structure. For values of the parameters, see Table S1. Setting 2 is identical to setting “H2” in [15] and the parameters used are identical.

| | $N$ | $g$ | $p$ | $q$ | $\rho$ | # effects | $\Psi_\theta$ structure |
| --- | --- | --- | --- | --- | --- | --- | --- |
| 1 | 180 | 30 | 500 | 1 | 0 | 5 | scalar |
| 2 | 180 | 30 | 500 | 1 | 0.6 | 5 | scalar |
| 3 | 250 | 50 | 1000 | 1 | 0 | 5 | scalar |
| 4 | 250 | 50 | 1000 | 1 | 0.6 | 5 | scalar |
| 5 | 250 | 50 | 1000 | 1 | 0 | 10 | scalar |
| 6 | 250 | 50 | 1000 | 1 | 0.6 | 10 | scalar |
| 7 | 250 | 50 | 1000 | 3 | 0 | 10 | scalar |
| 8 | 250 | 50 | 1000 | 3 | 0.6 | 10 | scalar |
| 9 | 250 | 50 | 1000 | 3 | 0 | 10 | diagonal |
| 10 | 250 | 50 | 1000 | 3 | 0.6 | 10 | diagonal |
| 11 | 250 | 50 | 1000 | 3 | 0 | 10 | unstructured |
| 12 | 250 | 50 | 1000 | 3 | 0.6 | 10 | unstructured |
| 13 | 250 | 50 | 1000 | 5 | 0 | 10 | scalar |
| 14 | 250 | 50 | 1000 | 5 | 0.6 | 10 | scalar |

#### Genome-Wide Association Studies (GWAS) Simulations

For the second study, we investigated a small-scale GWAS regression. Whereas a large-scale GWAS might contain hundreds of thousands of SNPs, we anticipate the CGD algorithm, even with the active set modification, only handling problems on the scale of several thousand SNPs. Using the `ggmix` R package [57], we generated 100 GWAS design matrices of size *N* = 250 by *p* = 1000 with entries representing minor allele counts in a diploid organism (0, 1, or 2). We partitioned the 250 samples from each design matrix into *g* = 10 clusters of size *n*_*i*_ = 25, significantly fewer than in the gene expression simulations (settings 3-14 of the gene expression simulations had *g* = 50 clusters, for example). This reproduces a more realistic GWAS design, with clusters representing different, perhaps geographically dispersed, populations. From each of these design matrices, we used the five random effect structures considered in the gene expression simulations (1 random effect, 3 random effects with scalar, diagonal and unstructured covariance matrices, and 5 random effects with a scalar covariance matrix) to generate five different response vectors *y* according to the mixed-effect model in Equation (1), with the same values of the fixed and random effect parameters as used in the gene expression simulations (Table S1).

#### Microbiome Simulations

Finally, we explored the application of high dimensional mixed-effects models fit by CGD to microbial operational taxonomic unit (OTU) data. OTU count data is compositional and characterized by heavy right-skew and high levels of sparsity [58]. To simulate microbiome data with realistic sparsity levels and covariance structures, we relied on functions in the R packages `SPRING` and `SpiecEasi` [59, 60]. At a high level, we generated our count data to mimic the marginal distributions of the 127 OTUs measured in the American Gut Project [61], and we specified different OTU correlation structures in our generated count matrices based on an assumed underlying microbe network. In particular, we considered the six different OTU-network structures provided in the R package `SpiecEasi` [59]: *band, cluster, scale free, Erdös-Rényi, hub*, and *block*. For each network type, we extracted the corresponding inter-OTU covariance structure and then generated 100 OTU count matrices following this covariance structure with sample size (number of rows) 120 and unique OTUs (number of columns) 127. The marginal distribution of the counts of a given OTU was forced to match the empirical cumulative distribution of the OTU counts from the American Gut Project data using the function `synthData_from_ecdf` in the R package `SPRING` [60]. Once we had generated these count matrices (100 for each network inter-OTU covariance structure, 600 in total), we transformed them by first adding pseudo counts to the zero entries and then applying a log-ratio transformation, using the last column as the reference. We partitioned the samples into 10 groups of 12 samples each, which are plausible cluster sizes for 16S sequencing studies.

From each simulated GWAS design matrix, we used the five random effect structures considered in the previous simulations to generate five different response vectors according to the mixed-effect model. To generate a response for a particular random effect structure with a particular normalized OTU design matrix, we first simulated a variable measured at the group level and appended it to the design matrix. We did this in order to study the ability of our model to handle group-level variables. We then constructed *Z* from the first *q* columns of this matrix, with *q* determined by the random effect structure under consideration. We finally generated the response *y* according to Equation (1). We retained the same fixed effect coefficients from the previous two settings, with the addition of a coefficient on the group-level variable chosen to be -1. However, we altered the random effect parameters for this setting in order to study how a different ratio of random effect to error variance might effect estimation performance (see Table S2).

### Applications to Real Omics Data

In addition to our simulations, we investigated the application of our implementation of high dimensional mixed-effects model on three published data set representing real examples of the three study types.

#### Bacterial Gene Expression and Riboflavin Production

The riboflavin data set is a popular high throughput transcriptomics data set with a longitudinal design [15]. Specifically, the data set contains repeated measures of the production of riboflavin inside cultures of bacteria (recombinant *Bacillus subtilis*) over a series of generations. Thus, observations are clustered within individual bacterial cultures with 28 distinct cultures, each having between 2 to 6 repeated measures at different time points, totaling 111 samples. For each observation, we have measurements of the expression levels of 4,088 bacterial genes and wish to use these to predict the riboflavin production rate.

#### Mouse GWAS Study for Body Mass Index

To demonstrate the ability of our method to fit larger-scale GWAS data sets, we applied it to data from a GWAS study in mice with 10,346 polymorphic markers measured in 1,814 individuals. This experiment was carried out by [62] to identify genetic signal for complex traits in a population of mice living in *g* = 523 different cages, which represented the grouping structure for our model. Using the high dimensional SNPs as predictors, we estimated a model for the body mass index (BMI) phenotype with cage-specific random intercepts. We included as additional unpenalized predictors in the model the age, gender and litter of the mouse (factor with 8 levels), imputing the median age for the 81 mice in which it was missing. This data was analyzed in a similar fashion by the authors of the `BGLR` package in R, using a Bayesian generalized linear model [53] that also included cage-specific random intercepts. Their model, however, did not include age as a predictor. This dataset was also re-analyzed recently in [50], who propose a quasi-likelihood estimation approach to fitting high dimensional mixed-effect models. Their model, however, does not include fixed litter effects. We comment on similarities and divergence between our results and these other analyses in the results.

#### Human Gut Microbiome Data across Age and Geography

Finally, we applied our model to a study of the diversity of microbial compositions across age and geography [63]. The data, available for download from Qiita (https://qiita.ucsd.edu/) with study ID 850, consists of microbial profiles (based on 16S gene sequencing) of individuals from three different countries ranging in age from infant to 83 years old. The original data set included 14,170 OTUs measured in 528 samples, representing 99 individuals from Malawi, 114 from Venezuela, and 315 from the USA. The inherent complexities of the actual data necessitated a meticulous approach to normalization, transformation, and data filtering prior to implementing our model. Following the pre-processing guidelines outlined in [28], we executed steps to eliminate less informative and noisy OTUs, along with employing transformation techniques to mitigate the impact of highly abundant taxa counts. Consequently, our data set underwent a refinement process, resulting in a subset of 1,362 OTUs for subsequent analyses. In [28], the authors apply a predictive model for age to this data, regularizing the OTU effects by assuming that they are components of a vector following a multivariate normal distribution with mean zero and an OTU-phylogeny-based covariance matrix. However, because their model could not accommodate the country-based clustering of samples, they included only the individuals from the USA in their analysis. In contrast, we model the age of individuals from all three countries using our penalized mixed-effect model, including country-unique random intercepts.

## Acknowledgements

This work was supported by the Department of Energy [DE-SC0021016 to CSL] and by the National Science Foundation [DEB-2144367 to CSL].

## Code availability

The high dimensional mixed-effects model is open source and publicly available as a Julia package `HighDimMixedModels.jl` in https://github.com/solislemuslab/HighDimMixedModels.jl. All simulation and real data analyses scripts are available in https://github.com/evangorstein/hdmmExperiments.

## Author contributions

CSL developed the initial idea for the study. EG and RA conducted all literature searches and performed all statistical analyses, including data preprocessing, model fitting, and summarizing results through figure creation. Additionally, EG developed the Julia package used in the study, while RA collected all the necessary data. EG drafted the initial version of the manuscript. RA and CSL contributed to interpreting the results and editing the manuscript. All authors have reviewed and approved the final version of the manuscript.

## Competing interests

The authors declare that they have no competing interests.

## Notes

### Competing Interest Statement

The authors have declared no competing interest.

